# Monitoring microscope performance in an imaging facility using OMERO-metrics

**DOI:** 10.64898/2026.06.28.735071

**Authors:** Sophia Sommer, Oumou Dhmine, Julio Mateos Langerak, Ian M. Dobbie

**Affiliations:** Baltimore Polytechnic Institute, 1400 West Cold Spring Lane, Baltimore, MD 21209, USA; Montpellier Ressources Imagerie, BioCampus, University of Montpellier, CNRS, INSERM, Montpellier, France; IGH, University of Montpellier, CNRS, Montpellier, France; Johns Hopkins University, Department of Biology, Mudd Hall, 3400N Charles St, Baltimore, MD 21218, USA

## Abstract

Microscopes are essential tools for discoveries on a scale invisible to the unaided human eye. The development of immuno-fluorescence followed by molecular biology techniques and fluorescent fusion proteins have revolutionised the use of optical microscopy in bioscience. The quality of the data produced is dependent upon the sample, its preparation and the instrument used. However, instruments can degrade over time without easily visible changes to the produced images and, in turn, negatively impacts results. By testing instruments and doing comparisons between results over time and between different instruments, problems can be highlighted and corrective action can be taken. Using small fluorescent beads the point spread function (PSF) of the microscope can be recorded and the image resolution measured. Beads were prepared in a concentration matched to the field of view size and dried onto coverslips and mounted on slides. The beads were then imaged as 3D Z-stacks of sufficient size to fully enclose the PSF of the system. This data was uploaded to OMERO and processed using OMERO-metrics, an OMERO plugin developed for this purpose. This paper summarizes the development of workflows and protocols to enable this process, presents the results obtained and demonstrates the detection of significant instrument issues.

## Introduction

Gathering scientific data requires rigorous understanding of the experiments being performed, including samples used, preparation of these samples and the instruments employed to study them. Due to setup errors, damage or just use over time, instruments can perform in a suboptimal manner, and a microscope is no different. However, with a microscope it is often not immediately obvious that the system is not performing at its best. With the difficulty of perceiving microscope performance issues, especially from images of arbitrary samples, system issues are often only flagged once they have become extreme, or after a specific adverse event. Slow degradation, or drift in performance are often not noticed and so remain uncorrected, producing less than optimal output data. These issues also affect reproducibility in experimental results^1^. The obvious answer to this is a rigorous and consistent set of tests, which are then analysed, allowing performance to be tracked over time and between instruments^2–5^. The importance of measuring and reporting such data has been stressed many times in the literature^6–11^. Although an ideal solution, this is very rarely performed on optical microscopes, due to the time and effort required to set up this type of workflow and then perform the actual steps including imaging, analysis and performance tracking. In a large facility such as the Integrated Imaging Center at Johns Hopkins University, where this work was performed, staff time is in high demand and such tasks are often given low priority meaning they are rarely, if ever performed. Streamlining the complete workflow is essential to enable recording and tracking these vital data.

There are several critical parameters that define the performance of a fluorescence microscope, including illumination intensity, illumination field uniformity, detection sensitivity and optical resolution^1^. In general, issues with illumination intensity, illumination uniformity or detection sensitivity are readily observed from normal usage, particularly on core facility based microscopes where many users are imaging samples they are very familiar with over time.

Users will notice issues with overall image uniformity or brightness and report these to facility staff who can investigate. However, changes in instrument resolution are often subtle and difficult to observe without careful measurement of well understood samples^12,13^. With this in mind, we set out to implement protocols and workflows to measure and track instrument resolution in a number of instruments and over time.

The broad approach is based on that of Nelson et. al.^14^. Standard slides of sub-resolution, 175 nm diameter, fluorescent beads are imaged, the resultant images analysed and the measured resolutions recorded and tracked between instruments and over time. It should be noted that the data in this paper presents Full Width Half Maximum (FWHM) values for Gaussians fitted to the imaged bead profiles. Although this differs from the theoretical definition of resolution^15^ and the commonly used Rayleigh Criterion^16^, it is easily measured, closely related and frequently used^3,5,17^. There are a number of other resolution measures such as Fourier Ring Correlation^18,19^, however, these are not as closely related to the theoretical definition of resolution or as easily applied to the standard bead samples we are using.

There have been several tools produced to automate analysis of such images often, summarising dataset to produce reports of current performance. Examples of these tools include MetroloJ/MetroloJ-QC^3^, PSFj^5^, as well as commercial packages such as Huygens (SVI, NL) or Daybook (Argolight, FR). Although extremely useful, these tools only automate a portion of the whole data workflow, not dealing with data organisation, result collation, or tracking results over time. Recently, the OMERO-metrics webapp for the OMERO^20^ image database system has been released (https://github.com/MontpellierRessourcesImagerie/omero-metrics) currently as a beta version. OMERO-metrics utilises OMERO to organise raw image data, automates the analysis and then tracks results over time providing easy to visualise image and graph output as well as data tables of numerical data.

In this study we have utilised the OMERO-metrics to measure and track the resolution performance of most of the fluorescence microscopes within a large imaging facility, the Integrated Imaging Center at The Johns Hopkins University Homewood campus. We have 13 fluorescence microscopes, most with several objectives, and multiple possible excitation and emission wavelength combinations. Ideally, the performance of all possible combinations would be tracked over time and compared between instruments and with theoretical values to ensure optimal performance. In this paper we present data from 9 of these systems including widefield, laser scanning and spinning disk confocal and light sheet instruments.

## Methods

We image the point spread function from a number of microscopes, fitted them with Gaussians and tracked data including the FWHM size X, Y and Z, asymmetry and quality of fit. To do this we used OMERO-metrics as this significantly simplified the workflow and reduced the effort required to systematically collect and analyse resolution data as measured by the PSF (Fig 1) and track it over time. The workflow involves capturing 3D images of sub-resolution beads and then uploading them to specific groups, projects and datasets on the OMERO server running the OMERO-metrics webapp. Once the data is uploaded each dataset simply requires confirmation of which images to include and review of the analysis parameters to start the automated processing, producing robust fits, individual bead data and data set averages, standard deviations and time courses of results, a diagram of the workflow is shown in Fig 1.

**Figure 1:**
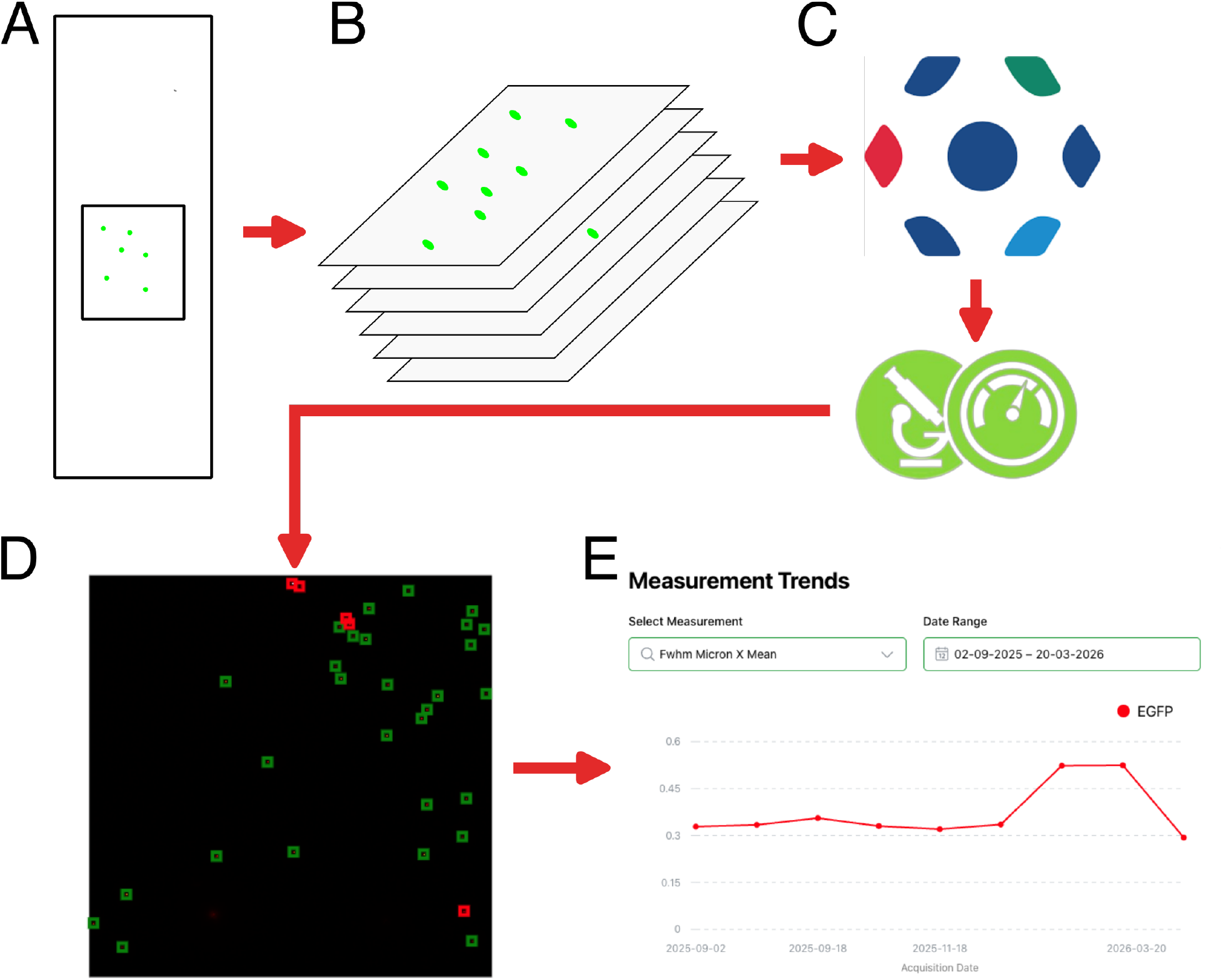
OMERO metrics workflow general approach diagram, A) A bead sample mounted on a slide, B) a 3D image stack is recorded from a microscope. C) the image data is imported into OMERO (top logo) and analysed with OMERO-metrics (bottom logo), D) An example analysed image stack within a dataset. A maximum intensity projection in Z with identified beads, green ROIs are valid beads, red ROIs were rejected for various reasons. E) The project level output can track metrics such as the FWHM in X across different datasets for a single instrument. Information flow is indicated by the red arrows.

We measured the resolution of most of our optical microscopes using standard 0.175 µm single colour fluorescent beads (Thermo Fisher, USA, p7220) which produce bright sub-resolution objects allowing relatively easy assessment of the 3D image resolution even on systems with lower magnification and NA, and hence lower sensitivity. These slides were imaged on a range of microscopes, usually with a 60x or 63x 1.4 NA oil immersion objectives. Mostly, 10 µm deep 3D stacks were recorded at between 0.2 and 0.3 µm Z slice spacing, exceptions for lower resolution systems are noted. When possible, we imaged 4 colours, named here as blue, green, red and farred, some systems did not have the correct excitation or emission wavelengths for all these wavelengths, see supplemental table 1 for instrument details, including excitation and emission wavelengths.

The systems available included 5 laser scanning confocals, 2 spinning disk confocals, 2 inverted widefield fluorescence systems, one upright fluorescence setup with a zoom body, one light sheet system, one lattice light sheet system and one conventional upright fluorescence microscope. Of the available systems we recorded data from 5 laser scanning confocals, 2 spinning disk confocals, one inverted widefield and one upright system with a zoom body, giving a total of 9 systems. The details of the system used for this paper are presented in Supplemental Material Table 1.

### Slide Preparation

We used 0.175 µm diameter Invitrogen p7220 PS-Speck beads with four different excitation/emission wavelengths: blue (350/440), green (505/515), red (540/560 called orange in the bead kit), and farred (633/660). Concentrated bead solutions were sonicated for 2 mins to split up aggregated beads then 10 μl of bead solution was added to 990 μl of ultra-pure water to produce a 1:100 dilution. This solution was then diluted by a further factor of 10 by putting 100 μl of 1:100 dilution into 900 μl of water to produce 1:1000 dilution. A further factor of 10 dilution was performed to get two solutions of 1:1,000 and 1:10,000. This process is repeated for each colour of beads. 200 μl of a dilution was then placed on to high precision coverslips, thickness 0.170 土0.005 mm (18 mm x 18 mm, Zeiss). The coverslips are left to dry overnight, in the dark, to allow the beads to stick to the coverslip surface. Finally, a drop of the provided mountant (part of the Invitrogen kit p7220, RI ∼ 1.47) is added to a slide and the coverslips inverted onto this drop, so the beads are in the mountant between the slide and the coverslip. Once surface tension has spread the mountant to the edges of the coverslip it is sealed with nail varnish and allowed to dry. These slides are kept in the fridge when not in use and last several months, although the blue beads tend to fade faster than the other wavelengths.

### Imaging and Analysis Protocol

1. Care was taken to ensure that data was sampled at sufficient spatial frequency to fully capture the optical resolution of the system. Systems were set for the relevant excitation and emission wavelengths. Z slice depth was set to the optimal as calculated by the software controlling that system, this takes account of both objective NA and the imaging wavelength. The laser scanning confocal systems were set with pinholes at 1AU and pixel sizes set to the optimum specified by the control software. System sensitivity was set so that the in focus beads filled more than half of the full dynamic range to maximise image signal-to-noise, this was done by adjusting the Illumination intensities, detector gains, exposure times and scan speeds depending on the exact controls present on each system. Experimental conditions could easily be repeated across image sessions by utilising the “Reuse” settings button present in most microscope control software.
2. Z stacks of 10 µm were recorded centred at the in-focus plane of the beads, except for the Zeiss Axiozoom, a zoom body system, where the minimum Z slice depth was 1 µm so that was used, and total stack height on this system was larger at 40-100 µm. Additionally on this system the zoom was set to maximum; this maximizes the NA of the objective utilised and hence resultant image resolution.
3. After collection, data was uploaded to an OMERO server using OMERO-insight client and stored in a group defined by the instrument, project defined by the objective and bead colour, and a dataset defined by the collection date.
4. OMERO-metrics analysis parameters were defined at the project level within the metrics view (Fig 2A), ie for each objective and bead colour, with the default parameters used for most settings except, SNR threshold=3, Fitting Gaussian R2 Threshold = 0.85 fitting Airy R2 threshold = 0.4, Bead Diameter Micron = 0.175. The bit depth was set depending on the image bit depth, required to ensure the saturation parameter is functional for instance, 12 bit data is saved as a 16 bit file.
5. Datasets were analysed once all images for a given dataset were uploaded. Note that analysing more data requires removing the previous analysis and reanalysing all images in a dataset. The analysis typically takes a minute or so per image stack in the dataset. This produces a summary output (top of Fig 2B) showing total number of beads found and fits deemed acceptable. Numerical outputs can be found by clicking on the dataset again while in metrics view (bottom of Fig 2B). The results will also then be accessible at the project level, with a summary over time of a given parameter and key values from a single dataset can be seen by clicking on a dataset in the measurement trends.

**Figure 2:**
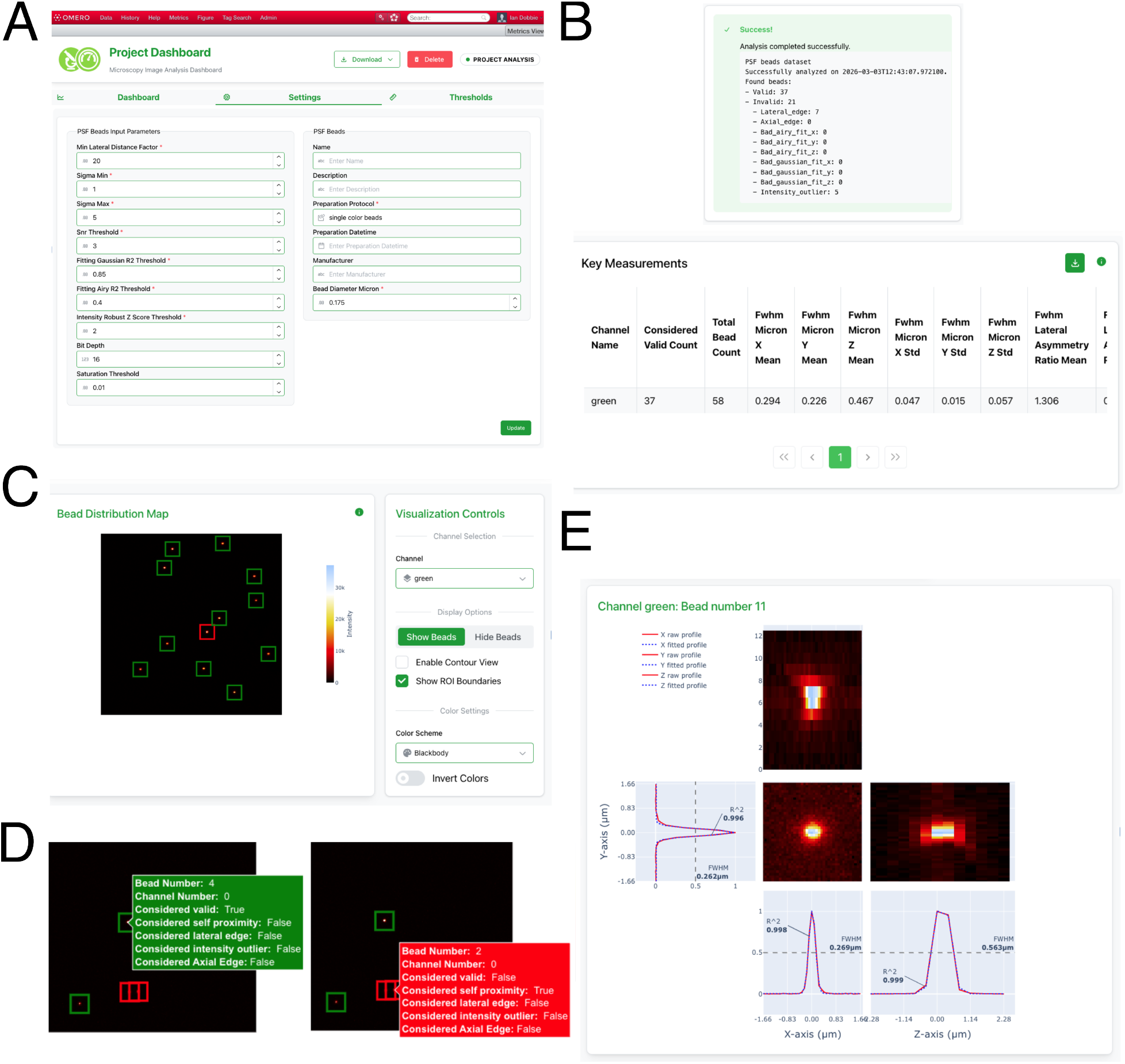
Typical PSF processing, A) Setting the analysis parameters for a specific project, B) The results screen and an example of some of the key values for a specific dataset. C) the metrics view of a single image stack, displaying the max intensity image and ROIs around identified beads. D) an example of an included bead with green ROI and displayed box and a rejected bead with red ROI and displayed box saying the bead was rejected as it was too close to another bead. E) The single bead metric view showing orthogonal views and profile plots along with fits and the fit quality (R^2^ value) and FWHM result for X, Y and Z, scale can be read from the profile axes.

Once the image data is processed the OMERO metrics view allows examination of the broad summary information of changes over time in the project level view, averages of various parameters in the dataset level view (Fig 2B, lower panel) and access to individual images (Fig 2C) and the ability to drill down to each individual beads.

The image level metrics view displays a maximum intensity image in Z along with boxed Regions Of Interest (ROIs) for each detected bead, green for accepted beads, red for rejected beads (Fig 2C). Hovering a mouse pointer over a green, or red, ROI displays information about the bead number and for a red ROI why it was rejected (Fig 2D). Clicking on a single bead will display orthogonal views of that specific bead along with the profiles, FWHM fit values and R^2^ fit parameters (Fig 2E). Additionally, the analysis stages also save an average bead image which, if selected in metrics view, will display orthogonal projections, profiles and fits of the average bead, similar to the single bead view. All the numerical data is also available via Attachments to the dataset which store information about every bead fit, reasons for rejection, profiles and other results. These attachments can be viewed as OMERO.tables directly in a web browser or downloaded as CSV or JSON files.

## Results

We used a standardised protocol (see Methods) for measuring single colour fluorescent bead samples across a range of microscopes. This allowed comparison of the performance of the systems against each other and over time to ensure that performance was stable.

The samples were all prepared in house from a single batch of beads to ensure that variability between samples was minimized. In general, we imaged at four different wavelengths, classified as blue, green, red and farred, again this varied slightly between systems as not all the microscopes had the ability to illuminate or image at these wavelengths. We imaged these samples on 5 laser scanning confocals, 2 spinning disk confocals, an inverted widefield, and an upright zoom body system.

The results from the point spread function measurements are presented in figures 3-6. We present lateral FWHM data in the X direction, but similar values were obtained for FWHM in Y (see supplementary material Fig S1). Most of the data was collected by a single user (SS) but a representative sample of systems and wavelengths were also collected by others, the different experimenters produced statistically indistinguishable results.

**Figure 3:**
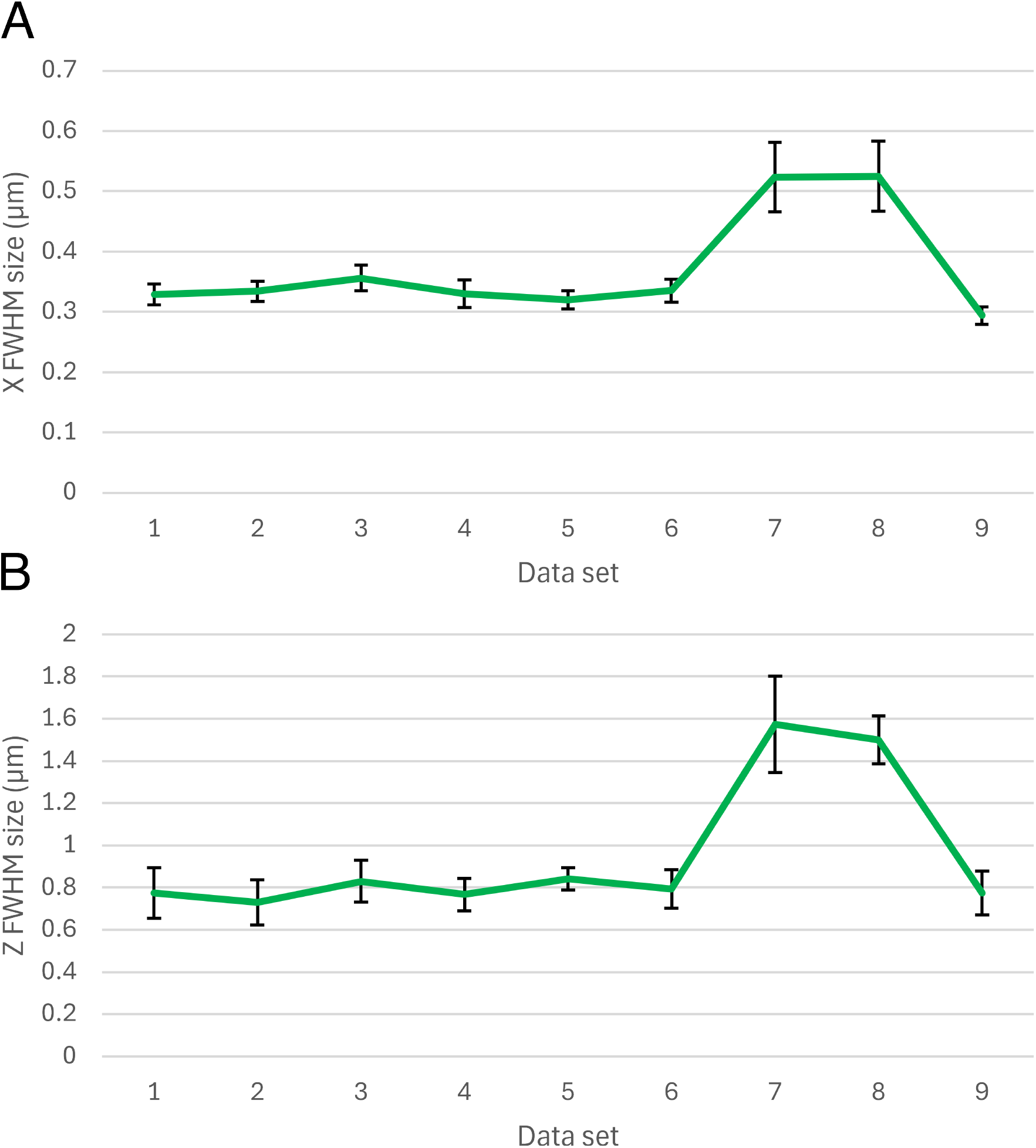
Tracking instrument performance over time. The datasets are from between September 2025 and March 2026. A) the FWHM of beads in the X direction from an Axio Observer widefield system, excitation at 480 nm, emission at 525 nm, note this is the same data as in Fig 1E but with error bars. B) the Z FWHM PSF size from the same data. Error bars are standard deviation with n=43,40,39,8,147,30, 55, 19 and 113 in both panels.

We cover four comparative data sets in this paper, demonstrating a range of uses for this streamlined workflow to allow testing several systems, including repeat testing. These data sets cover:

1. Following the performance of a single system over many testing sessions, including data collected by different users to demonstrate consistency and repeatability.
2. Comparisons of data from a single instrument with two different objectives, one of which has been damaged and shows a significantly aberrated PSF shape and increases in the FWHM in lateral and axial directions.
3. Comparing the same instrument across imaging modes, showing widefield, conventional spinning disk confocal and SoRa super-resolution spinning disk confocal results.
4. Representative data from 9 instruments allowing performance comparison between these instruments.

### Tracking performance over time

We followed the performance of an inverted widefield system using green beads and a 63x 1.4 NA oil immersion lens. Measurements were taken by 4 different people over the period from September 2025 to March 2026, the FHWM of the bead images were used to track the instrument resolution, Fig 3. It can be seen that the resolution is consistently ∼330 nm in X (mean 334 nm ± 12 nm SD datasets 1-6) and ∼790 nm in Z (mean 790 nm ± 42 nm SD datasets 1-6) until measurement 7 where the measurements become significantly worse at 525 nm in X and >1500 nm in Z. We recorded a second data set on the same day with the DIC prism removed to see if the issue was related to either the prism itself, or to dust/dirt on the prism.

This had no significant effect on the results, compare datasets 7 and 8 in Fig 3. The system was then thoroughly cleaned, including the objective and camera. No obvious issues were seen during this process, although the camera did have noticeable dust on its front glass surface. The camera dust would not cause global resolution loss as its effect would be localized to regions of the image near the dust, as the surface is close to the image plane on the camera chip. The low-resolution measurements are from 74 beads (55 with DIC prism and 19 without) randomly spread across the camera in 11 total image stacks. After cleaning, dataset 9 in Fig 3, produced resolutions very similar to those recorded earlier in datasets 1-6.

### Comparative study of a damaged and an undamaged objective

We did a comparative study of two objectives on one system after one of the objectives suffered accidental damage. The system was a Zeiss LSM 800 laser scanning confocal with two nominally identical 63x 1.4 NA Plan-Apochromat oil immersion objectives. The results from this comparison showed that the XY and Z resolutions were significantly worse with the damaged objective compared to the undamaged one. A distinct curve shape and elongated Z size can clearly be seen in the orthogonal projections of a single bead shown in Fig 4A compared to the more symmetrical Fig 4B. The decrease in the FWHM in X for all 4 colours in the undamaged objective can be seen in Fig 4C.

**Figure 4:**
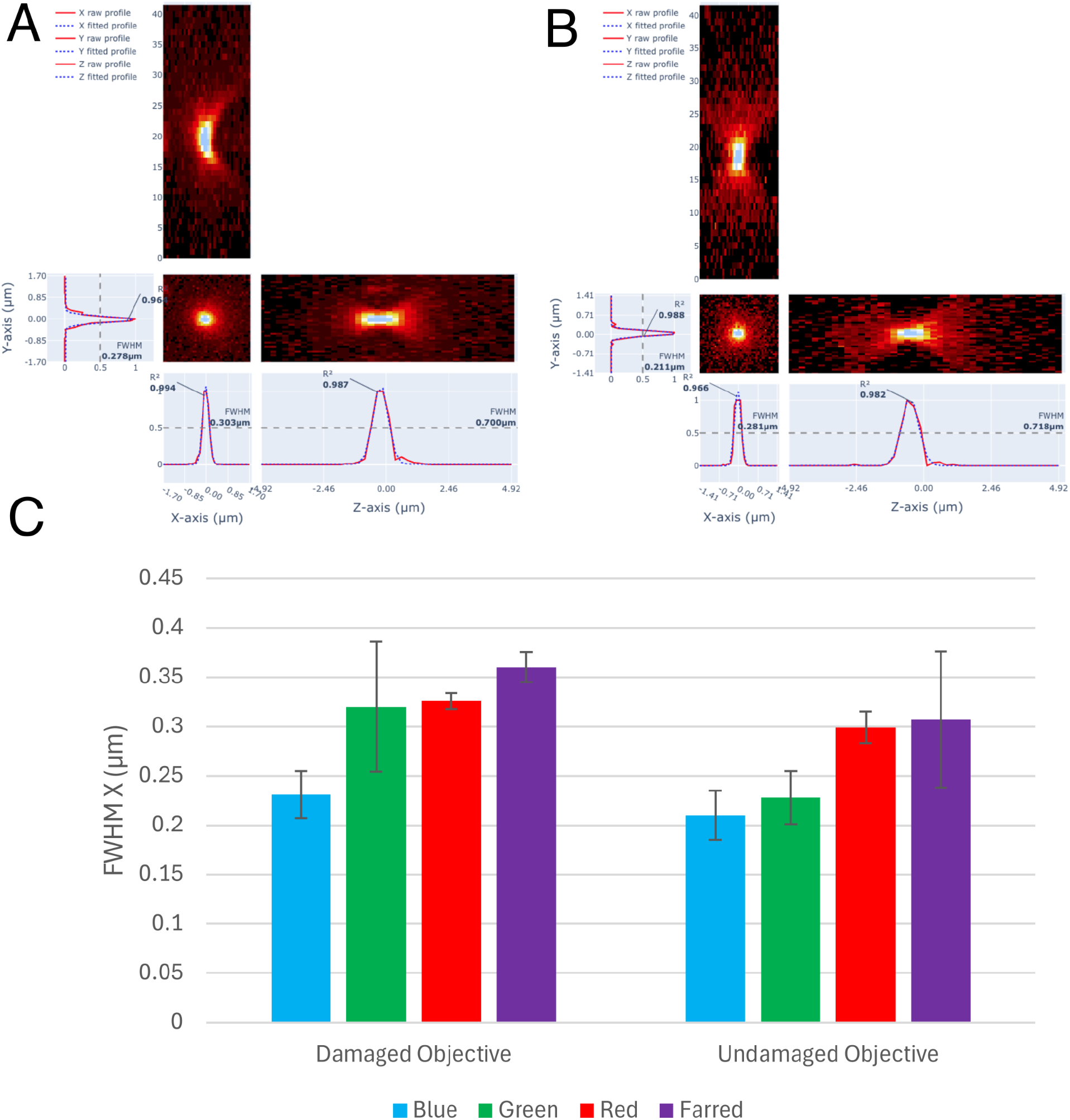
Images of a single bead at 488 nm excitation and 520 nm emission from a Zeiss LSM 800 confocal with the pinhole set to 1AU, A) with a damaged objective, showing XZ, XY and YZ projections as well as profiles along the Y, X and Z axis including the raw data and fitted Gaussians. Fits include an R^2^ fit quality measure and the calculated FWHM. B) The same information with an undamaged objective. C) Lateral FWHM in X showing the degraded resolution from the damaged objective (N = 55,57,34,8,17,30,15 and 16 from left to right).

### Different imaging techniques on a single system

As an additional comparative study, we used all 4 colour beads to compare the resolution achieved on a single system in widefield, with a conventional spinning disk confocal unit and with a SoRa super resolution spinning disk module^21^. These images were taken on a Nikon system with a 60X 1.42NA objective for the widefield and conventional spinning disk images and a 60x 1.49NA TIRF objective for the SoRa disk images. The Super resolution images also included an additional 2.8x magnifier in the optics to reduce the projected pixel size to ∼40 nm to properly sample the enhanced resolution. The resolution was very similar between the widefield and conventional confocal images, Fig 5. The resolution is then significantly improved by the SoRa confocal spinning disk module. The SoRa resolution is above the theoretical value with the green bead FWHM measured at ∼225 nm. This is almost the expected resolution improvement of 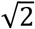, but significantly worse resolution than the expected ∼150 nm. The beads used, at 175 nm diameter, are no longer a sub resolution object at this resolution which will contribute to the higher FWHM than might be theoretically expected.

**Figure 5:**
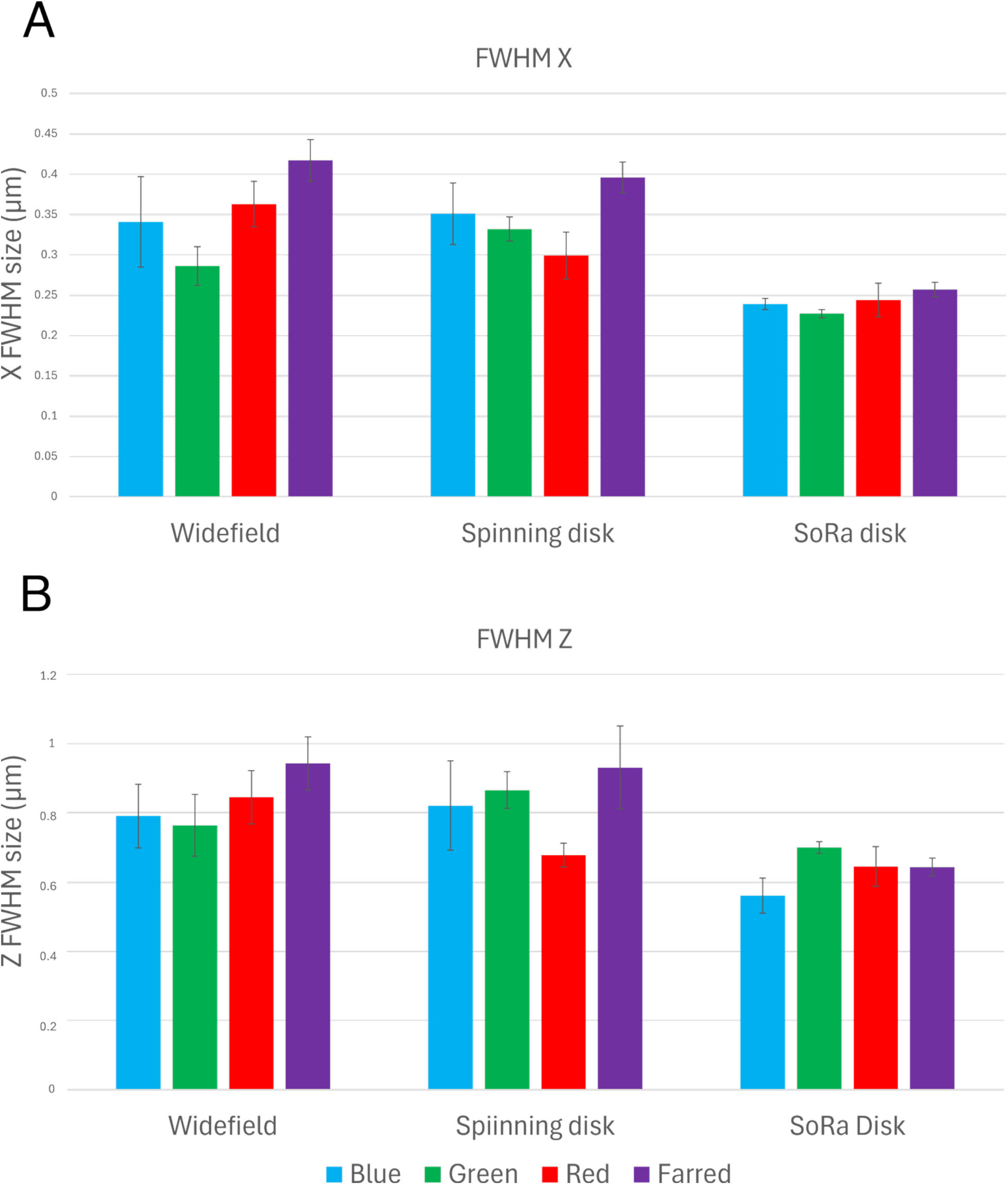
Comparison between widefield, conventional spinning disk confocal and a SoRa super-resolution spinning disk confocal all on the same system. A) Average FWHM in X for the 3 modalities in blue, green, red and farred, B) as A but for FWHM in the Z direction. Note that the widefield and conventional spinning disk data were taken with 1.42 NA objective while the SoRa data came from a 1.49 NA TIRF objective. N=34,47,107,123,27,38,61,34,11,20,71 and 31 left to right in both A and B.

### Comparison across many systems

Finally, we gathered data from all of the systems we measured (Fig 6). It is clear that the FWHM values trend up with wavelength as expected, with a few outliers, eg the red values on the LSM700-17 and green on the LSM980, this could be due to statistical variation over the 33 measured values presented. The values fall in the range 0.2-0.4 µm range expected for optical resolution in systems with an NA of 1.4, with two exceptions. The Zeiss SD system which seems to be substantially worse than expected, with a 1.4 NA lens, and the Axiozoom system which has a much lower NA at 0.57. The limited NA of the Axiozoom system is particularly evident in the Z FWHM data (Fig 6B) as it is many times worse than the other systems, at 5-8 µm compared to < 1 µm. The lower resolution of the Axiozoom system is expected but the results are even lower than would be expected from the purported NA of the system. We suspect that the zoom body limits the real NA to approximately 0.3, and that both XY and the Z measured resolutions are worse than expected as a result.

**Figure 6:**
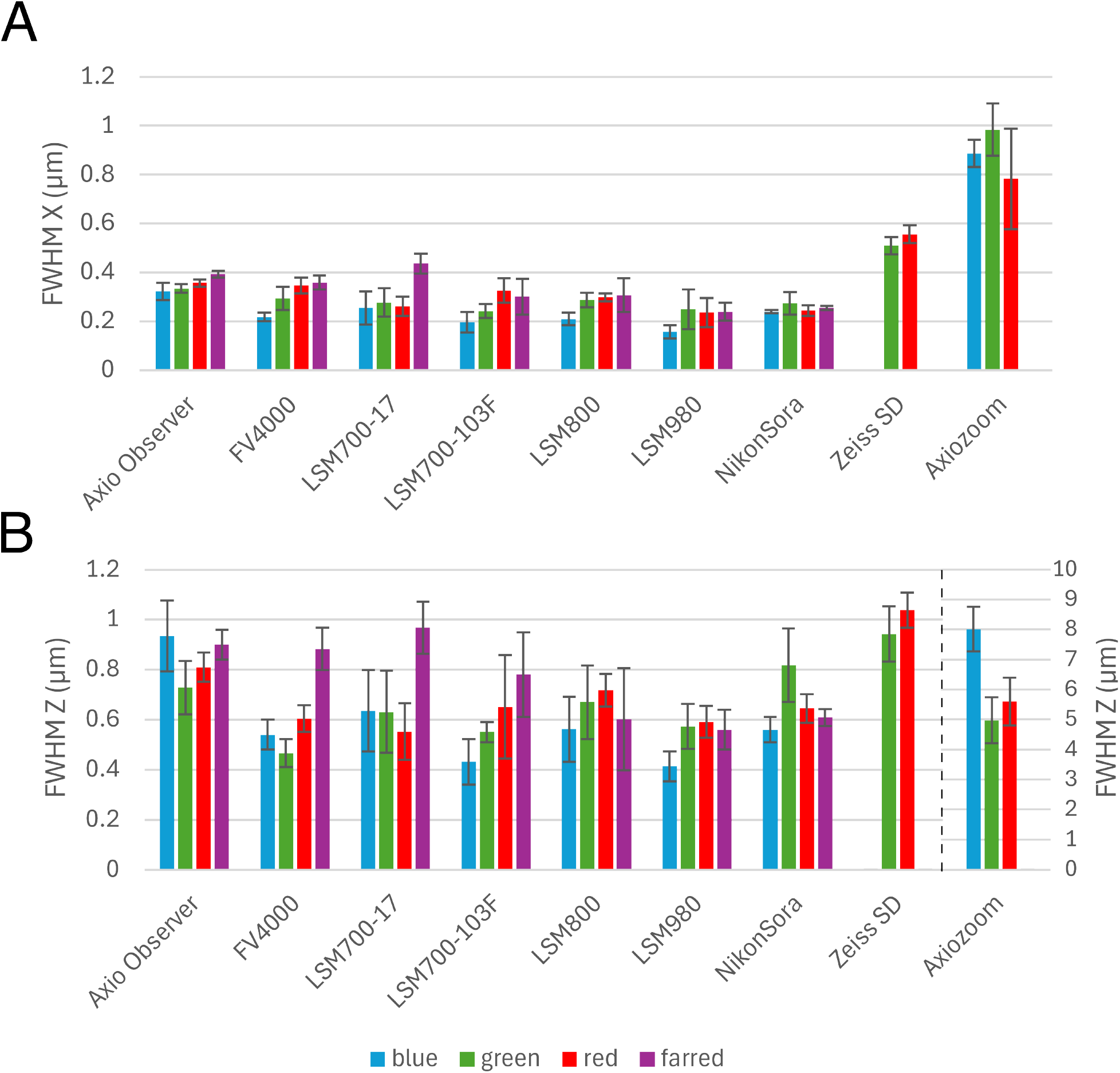
Comparison of FWHM PSF size between 9 instruments. A) FWHM PSF size in X as derived from a Gaussian fit of a 3D image of a 175 nm bead. B) the same instruments and bead images but now in the Z direction, with the Axiozoom data on the secondary Y axis on the right. N=40, 40, 89, 81, 50, 37, 51, 44, 9, 18, 29, 8, 40, 18, 32, 14, 55, 15, 34, 8, 15, 31, 21, 6, 11, 29, 71, 161, 37, 15, 10, 41 and 60 from left to right.

The Zeiss SD system is obviously underperforming and we investigated the cause. The system was set up in such a way that the disk pinholes were not properly focused on the camera image plane. When this was corrected the lateral resolution significantly improved to ∼375 nm in green and ∼ 390 nm in red (see supplemental data, Fig S2). This is still significantly worse than might be expected but the system has EMCCD cameras with relatively large pixels and the projected pixel size is 210 nm leading to a significant degradation in the produced image resolution due to the under sampling of the optical resolution.

## Discussion

We have shown that OMERO and OMERO-metrics is a very effective tool to monitor microscope resolution performance via standardised imaging protocols, automated analysis and result tracking. The measurement we performed clearly showed deficiencies in 3 systems out of 9 studied, including damage caused by an accident on one system, a critical setup error on a second system and the reduction in resolution of a third system over time, likely due to dirty optics. Finding these issues allowed us to address them and, except for the reported accident, it is unlikely we would have discovered the issues until later if at all.

We have developed a standard protocol that works well for our imaging facility, enabling regular assessment of resolution performance over a number of instruments. Additionally, over the period of this research we have worked to improve the user interface and core functionality of OMERO-metrics, with substantial improvements in several areas including curve fitting performance, access to important results, error reporting and increased speed of access to results down to the individual bead level. This is an ongoing process with the tool continuing to improve. This work focused on the PSF measuring module, but the software can also measure and track field flatness. A workflow and analysis routines for laser power stability over both short and long term time scales is currently being developed. The core developers of OMERO-metrics will publish a paper on the details of the software in future.

OMERO-metrics provides a convenient, fast and easy to use mechanism to take collected PSF images, analyse them and then collect both the input data and the results for comparison. We found that system to be robust and reliable, with multiple measurements of the same system to produce similar results over time and with both the same or different operators. In fact over the roughly 9 month period of data collection, we only saw significant changes in the performance of one instrument. However, this did indicate a degradation of about 60% in the XY performance and almost 100% in Z. Investigation pinpointed no obvious cause, however thorough cleaning of that instrument returned it to the previous performance levels (Fig 3) showing the usefulness of this data collection and analysis strategy.

The combination of developing a simple workflow and utilising automated analysis and data tracking within OMERO-metrics has significantly improved the information available about our microscope systems. This enabled the identification of one system that is significantly underperforming, the Zeiss SD system, and significant degradation in another system that was quickly identified and corrected with minimal impact on users of the imaging facility. Overall, OMERO-metrics significantly lowers the barriers to implementing regular testing of resolution measurements across systems improving reproducibility and traceability of results from these systems.

### Contributions

The project was conceived, planned and run by IMD. Almost all data collection and analysis were performed by SS under the guidance of IMD. OD and JML developed OMERO-metrics and made substantial alterations to enable and improve the data analysis, error reporting and UI for this project. This paper was written by all authors.

## Acknowledgements

The majority of the work in this paper was performed by a high school student enrolled on the Ingenuity Project (https://www.ingenuityproject.org/) at Baltimore Polytechnic Institute. This work would not have been possible without the effort of the QUAREP (https://quarep.org/) community pushing forwards the expectation that instrument performance is a vital component of the imaging process and that measuring this performance should be both easy and routine.

A small amount of image data was collected by graduate students at Johns Hopkins University taking a graduate course in Optical Microscopy (AS.020.608 Autumn 2025).

We thank France BioImaging (https://france-bioimaging.org/) for funding the salary of OD (supported by ANR-24-INBS-0005 FBI BIOGEN).

This work would not have been possible without the support of the staff at the Integrated Imaging Center, JHU, particularly Erin Pryce and Yuan Cai.

Grant support: NIH grants #1S10OD010712-01A1, #1S10OD020152-01A1, #1S10RR019409-01, #1S10 OD021567-01A1.

