## Supplemental Tables and Figures for "Monitoring microscope performance in an imaging facility using OMERO-metrics"

Table 1 - details of imaging systems

| Name | Microscope stand | inv/upright | modality | objective | light source | detector | excitation blue | excitation green | excitation red | excitation farred | emission blue | emission green | emission red | emission farred |
|---|---|---|---|---|---|---|---|---|---|---|---|---|---|---|
| Axio Observer | Zeiss AxioObserver 200M | inverted | WF | 63x 1.4 oil | Metal halide | Flash 4.0 | G365 | 470/40 | 550/25 | 640/30 | 445/50 | 525/50 | 605/70 | 690/50 |
| FV4000 | Evident FV4000 | inverted | LSM | 60x 1.42 oil | Laser | SilVIR | 405 | 488 | 561 | 640 | 430-470 | 500-550 | 570-620 | 650-710 |
| LSM700-17 | Zeiss AxioObserver Z.1 | inverted | LSM | 63x 1.4 oil | Laser | Alkali PMT | 405 | 488 | 555 | 639 | 300-584 | 493-800 | 568-800 | 300-800 |
| LSM 700-103F | Zeiss AxioObserver Z.1 | inverted | LSM | 63x 1.4 oil | Laser | Alkali PMT | 405 | 488 | 555 | 639 | 300-584 | 493-800 | 568-800 | 300-800 |
| LSM 800 | Zeiss AxioObserver Z.1 | inverted | LSM | 63x 1.4 oil | Laser | Alkali PMT | 405 | 488 | 561 | 640 | 400-555 | 400-650 | 450-700 | 645-700 |
| LSM 980 | Zeiss AxioObserver Z.1 | inverted | LSM | 63x 1.4 oil | Laser | GAsP PMT | 405 | 488 | 561 | 639 | 410-553 | 491-677 | 562-695 | 642-695 |
| NikonSora | Nikon Eclipse Ti2 | inverted | WF | 60x1.42 oil | LED | Fusion BT | 385 (375/28) | 475 (480/30) | 550 (540/25) | 621 (620/50) | 460/50 | 535/20 | 605/55 | 690/50 |
| NikonSora | Nikon Eclipse Ti2 | inverted | SD | 60x 1.42 oil | Laser | Fusion BT | 405 | 488 | 577 | 640 | 445/49 | 531/50 | 647/100 | 691/64 |
| NikonSora | Nikon Eclipse Ti2 | inverted | SoRa SD | 60x 1.49 TIRF oil | Laser | Fusion BT | 405 | 488 | 577 | 640 | 445/49 | 531/50 | 647/100 | 691/64 |
| Zeiss SD | Zeiss AxioObserver Z.1 | inverted | SD | 63x 1.4 oil | Laser | Evolve | | 488 | 561 | | | 520/35 | 629/62 | |
| AxioZoom | Zeiss zoombody V16 | upright | WF | 2.13x 0.57 | Metal halide | MRm | G365 | 470/40 | 550/25 | | 445/50 | 525/50 | 605/70 | |

Table 1 Notes:
**Modalities**: WF - Widefield, LSM - Confocal Laser Scanning Microscope, SD - Spinning Disk Confocal.

**Detectors:** Flash 4.0 - Hamamatsu sCMOS v2, SilVIR - Evident Broadband for blue and green and Red-shift type for red and farred, Alkali PMT - standard Zeiss Alkali PMT, GAsP PMT - Zeiss GAsP PMT, Fusion BT - Hamamatsu Orca Fusion BT, Evolve - Photometrics Evolve Back illuminated EMCCD, MRm - Zeiss MRm CMOS.
**Excitation:** Laser wavelengths as a single number, Metal halide sources modulated by bandpass filters shown as center/width wavelengths in nm, LED sources shown as center LED wavelength with bandpass filter in brackets afterwards.
**Emission:** bandpass emission filters expressed as center/width, except G365 which has peak transmission at 365 nm and a width of about 50 nm, spectral confocal detectors shown as min-max wavelength. Note that these systems also include a dichroic and often a laser blocking notch filter, so the excitation wavelength is sometimes included in detection range.

### Comparison of FWHM in X and Y directions

Although FWHM measurements are not exactly the same in X and Y, there is very little difference between them as shown by data from the AxioObserver system. The X and Y values are presented in Figure S1 for the data sets 1-6 as shown in Figure 3 of the main paper, where the resolution is reproducible across a number of measurements. These 6 data sets were taken by 4 different experimenters over a month between September and October 2025. With this similarity and reproducibility, we only present lateral FWHM in X in the main paper to simplify the figures.

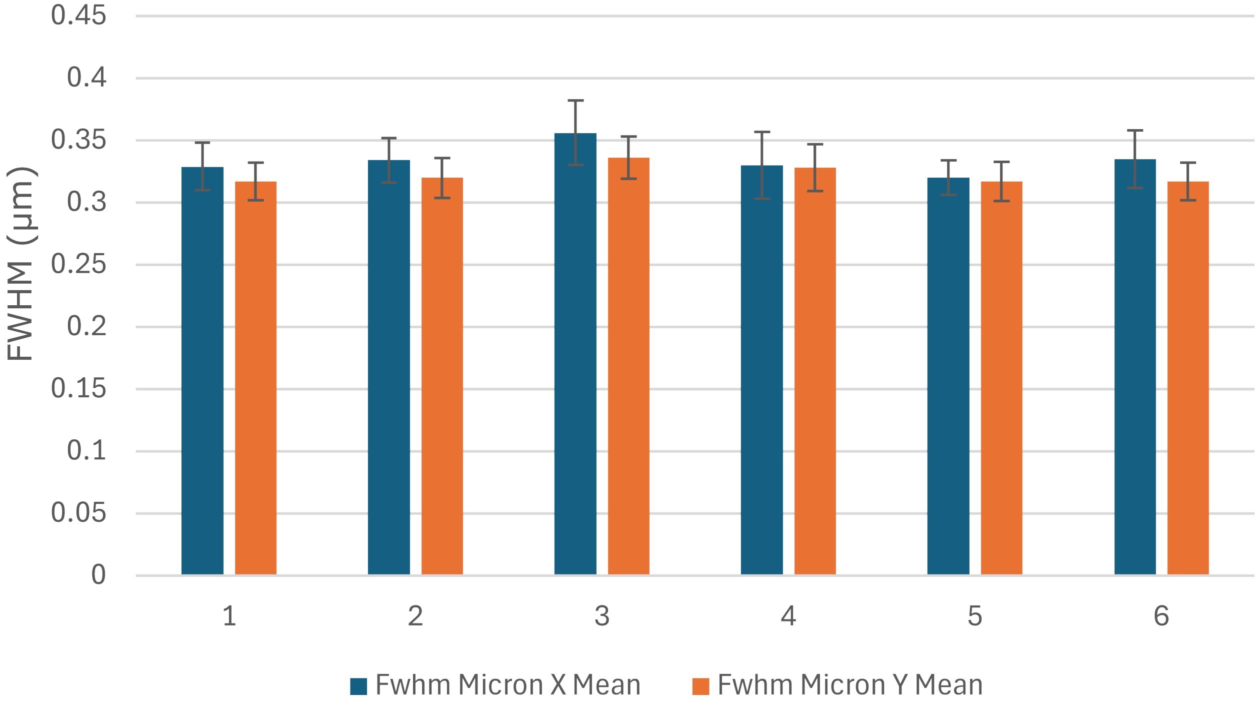


Figure S1: Comparison of FWHM measurements in X and Y for the AxioObserver system in green over data sets 1-6 as shown in figure 3. Error bars are SD, with N = 43,40,39,8,147 and 30 for the pairs of bars, each pair of bars has the same number of measurements as they are from the same set of beads.

### Comparison of the Zeiss Spinning Disk confocal performance

The unexpected large size of the PSF from the Zeiss SD system led us to investigate the system in detail. The culprit turned out to be misalignment in the system such that the disk pinholes were not properly focused on the camera. We refocused the camera, so it was focused on the disk pinholes and the FWHM significantly improved in X, Y and Z. The X and Z data for green and red beads are shown in Figure S2. The X measurement in green improved from 508±36  nm to 376±16 nm, so the PSF was reduced roughly 25% linearly in the green and was actually improved even more in the red. Overall the total volume of the PSF was reduced to less than 40% of its original size, so the reduction is isotropic, $0.75^3$ =0.42.

However, it should be noted that the 376 nm lateral FWHM is still substantially more than the expected resolution in the green with a 63x 1.4 NA lens. This is caused by the large pixel size of the EMCCD camera used producing a projected pixel size of 212 nm in the sample plane.  This is significantly larger than the required Nyquist sampling which would be about 100 nm. The undersampling leads to a larger than expected PSF and it can be seen that there are very few points across the peaks in X and Y in the profile graphs in Fig S2 B. Nyquist sample is typically achieved in our other systems which have CMOS cameras with 6.5 um pixels and a 60x or 63x objective, to give less than 110 nm projected pixel size in the sample plane. The laser scanning confocal systems have extensive control over the pixel size by electronically varying the total scan angle controlled via the zoom parameter in the software interfaces. This enables them to closely match Nyquist sampling over the whole range of objectives and wavelengths.

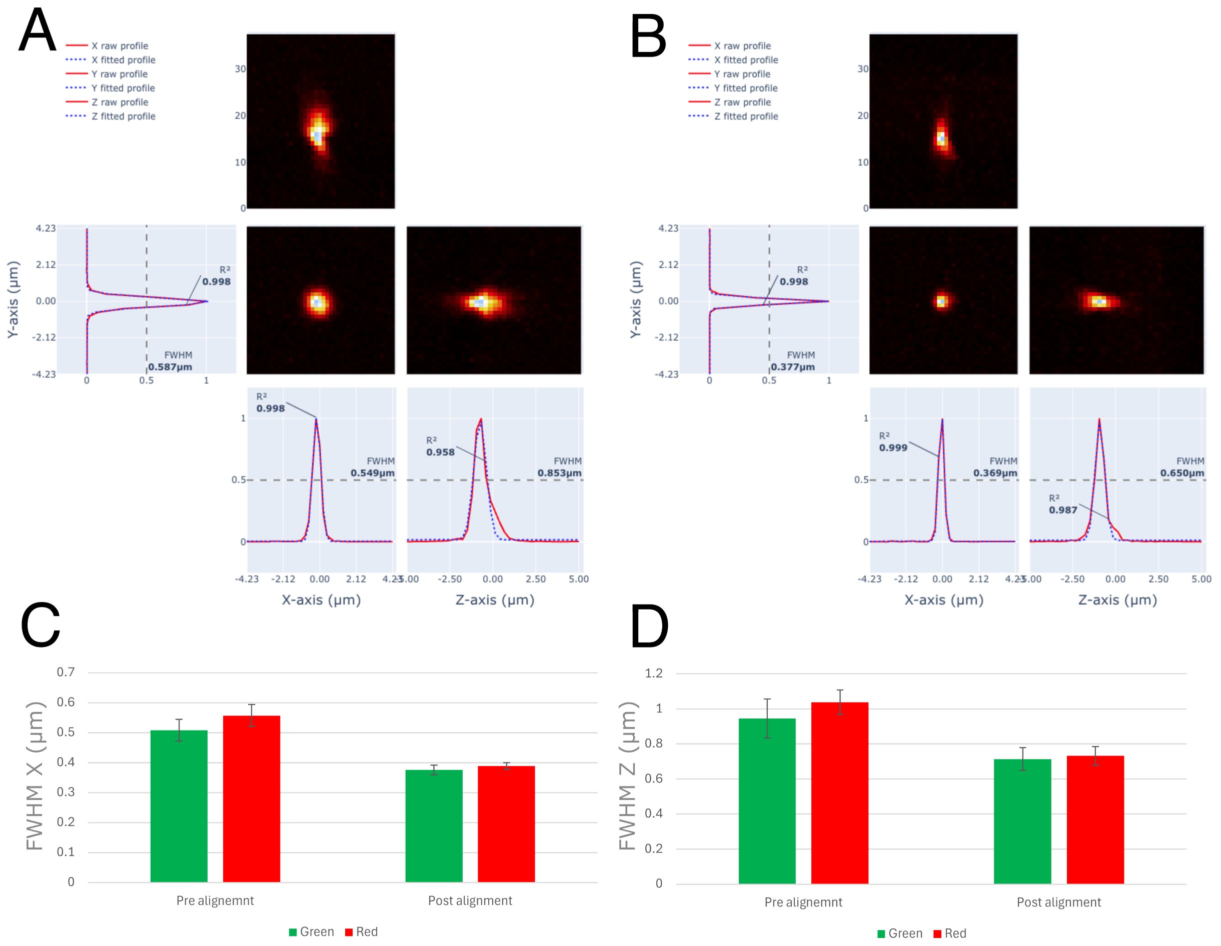


Figure S2: Comparison of Zeiss SD orthogonal projections and FWHM measurements before and after refocusing the disk pinholes into the camera image plane. A & B, orthogonal projections, profiles and fits of representative single green beads from pre- and post-alignment data sets. C & D FWHM mean values in X and Z for green and red beads, error bars are SD, N= 38,15,20 and 35 from left to right in each graph.
